# A hybrid cyt *c* maturation system enhances the bioelectrical performance of engineered *Escherichia coli* by improving the rate-limiting step

**DOI:** 10.1101/2020.04.01.003798

**Authors:** Lin Su, Tatsuya Fukushima, Caroline M. Ajo-Franklin

## Abstract

Bioelectronic devices can use electron flux to enable communication between biotic components and abiotic electrodes. We have modified *Escherichia coli* to electrically interact with electrodes by expressing the cytochrome *c* from *Shewanella oneidensis* MR-1. However, we observe inefficient electrical performance, which we hypothesize is due to the limited compatibility of the *E. coli* cytochrome *c* maturation (Ccm) systems with MR-1 cytochrome *c*. Here we test whether the bioelectronic performance of *E. coli* can be improved by constructing hybrid Ccm systems containing protein domains from both *E. coli* and *S. oneidensis* MR-1. The hybrid CcmH increased cytochrome *c* expression by increasing the abundance of CymA 60%, while only slightly changing the abundance of the other cytochromes *c*. Electrochemical measurements showed that the overall current from the hybrid *ccm* strain increased 121% relative to the wildtype *ccm* strain, with an electron flux per cell of 12.3 ± 0.3 fA·cell^-1^. Additionally, the hybrid *ccm* strain doubled its electrical response with the addition of exogenous flavin, and quantitative analysis of this demonstrates CymA is the rate-limiting step in this electron conduit. These results demonstrate that this hybrid Ccm system can enhance the bioelectrical performance of the cyt *c* expressing *E. coli*, allowing the construction of more efficient bioelectronic devices.

## 1. INTRODUCTION

The flow of electrons can transfer both energy and information between living and non-living systems for bioelectronic and biosensing applications. Extracellular electron transfer (EET) pathways (Shi et al., 2016) can electrically connect living microorganisms with electrochemical cells, enabling applications such as power generation in microbial fuel cells (MFCs) (Gul and Ahmad, 2019; Logan, 2009; Santoro et al., 2017; Slate et al., 2019) and environmental sensing in bioelectronic sensing systems (Chouler et al., 2018; Wang et al., 2013; Zhou et al., 2017). Engineering EET in native electroactive microorganisms, such as *Shewanella oneidensis* MR-1 (Cheng et al., 2020; Min et al., 2017; West et al., 2017), and non-native hosts, such as *Escherichia coli* (Jensen et al., 2016; Sturm-Richter et al., 2015; Thirumurthy and Jones, 2020) or *Pseudomonas putida* (Schmitz et al., 2015), has expanded the range of potential applications (Harris et al., 2017; Su and Ajo-Franklin, 2019). However, a fundamental limitation in these bioelectronic systems is the low rate of EET, which causes these devices to have low power output, low sensing sensitivity, and limited robustness. For example, expressing the EET pathway from *Shewanella oneidensis* MR-1 into engineered *E. coli* only yields ∼10% the current produced by MR-1 (TerAvest et al., 2014).

The molecular structure of these EET pathways is increasingly well-understood. The direct EET pathways (Mtr pathway) in *Shewanella oneidensis* MR-1 is a group of multiheme cytochrome *c* (cyt *c*) proteins embedded in the inner membrane (CymA), in the periplasm (Stc, FccA), and outer membrane (MtrC, MtrA) (Clarke et al., 2011; Okamoto et al., 2011; Rosenbaum et al., 2012). We and others (Jensen et al., 2016; Su et al., 2020; TerAvest et al., 2014) have found that expressing the CymA, MtrC, MtrB and MtrA proteins of the Mtr pathway enables *E. coli* to perform EET (**Figure 1A**).

**Figure 1.**
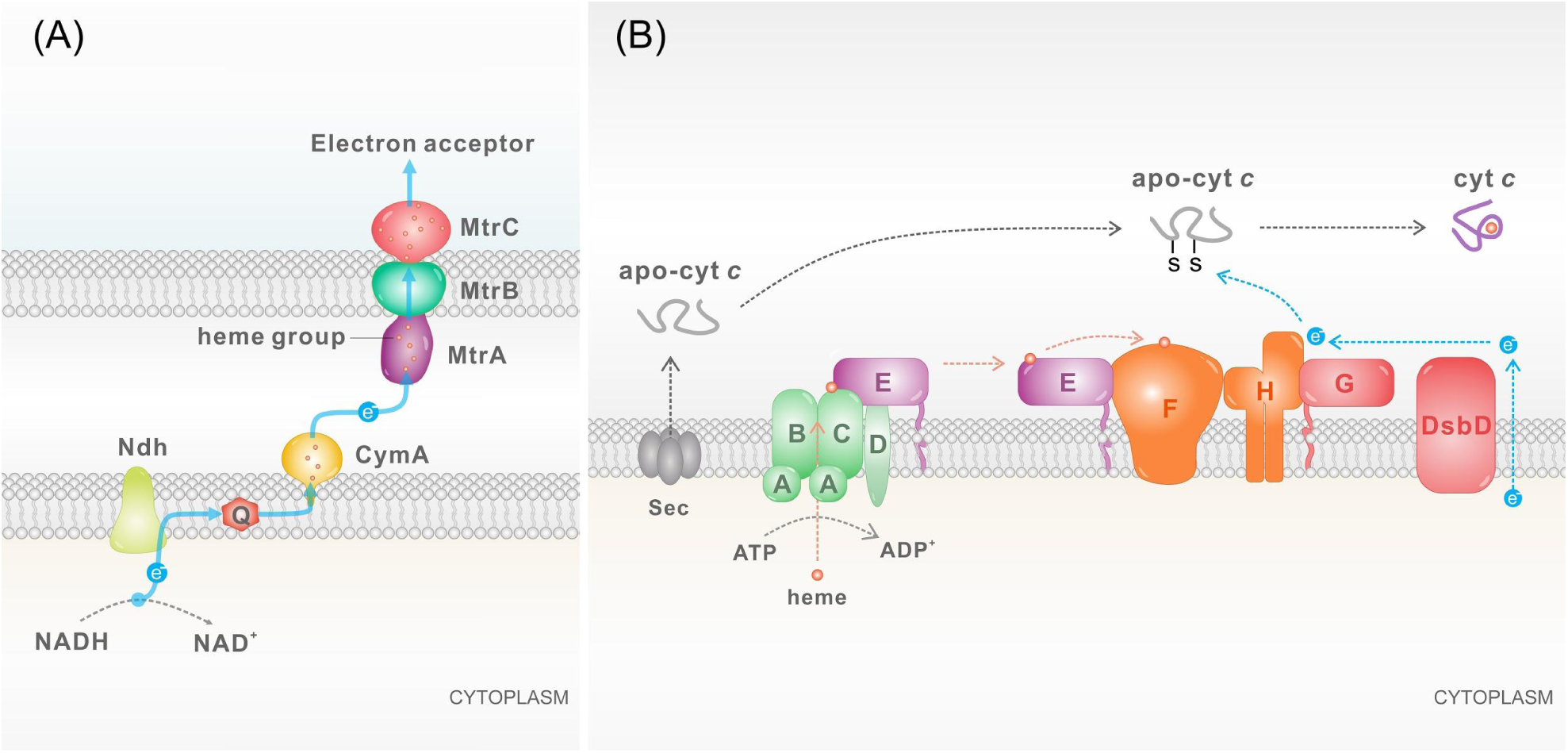
A schematic of the Mtr pathway in engineered *E. coli* and the three modules of Ccm system in *E. coli*. (A) The Mtr pathway includes MtrC, MtrB, MtrA and CymA from *Shewanella oneidensis* MR-1, which enable engineered *E. coli* to perform extracellular electron transfer. (B) For cytochrome *c* (cyt *c*) such as MtrC, MtrA and CymA to mature, the Ccm system catalyzes ligation of heme with apo-cyt *c* to form covalent bonds with heme groups. In *E. coli*, the Ccm system is composed of eight different Ccm proteins, CcmA, B, C, D, E, F, G, H, and a DsbD (disulfide bond formation protein). These proteins form three functional modules: heme delivery (green and purple), apo cyt *c* chaperone (red), and ligation (orange).

Expressing the Mtr cyt *c* proteins requires the cytochrome *c* maturation (Ccm) system, which covalently ligates heme *b* groups to the apo-cyt *c* protein to form the holo-cyt *c* (Sanders et al., 2010; Verissimo et al., 2011). The Ccm systems of α- and β-proteobacteria contains three modules (**Figure 1B**): the heme *b* translocation module that conveys heme *b* into the periplasm and to the ligation module, the apo cyt *c* chaperone module that maintains apocyt *c* in the reduced state, and the ligation module that catalyzes formation of a thioether bond between the heme b and apocyt *c* (Kranz et al., 2009; Sanders et al., 2010; Thöny-Meyer, 2000). In both *S. oneidensis* and *E. coli*, CcmA, B, C, D and E form the heme delivery module, and CcmG and DsbD compose the chaperone module. While *E. coli* uses two proteins, CcmF and CcmH, as the ligation module, this module in *S. oneidensis* MR-1 includes a third protein - CcmI (**Figure S1**). We recently found that mutations in the C-terminal domain of CcmH can significantly change the EET efficiency by changing the stoichiometry of the Mtr cyt *c (Su et al., 2020)*. This finding suggested that optimizing the heme ligation process by redesigning CcmH of *E. coli* could remodel the stoichiometry of cyt *c* and enhance the bioelectrical performance of Mtr-expressing *E. coli*.

The role of the C-terminal domain of CcmH in cyt *c* maturation is disputed. Fabianek et al. (Fabianek et al., 1999) and Robertson et al. (Robertson et al., 2008) demonstrated that *E. coli* strains with an incomplete CcmH lacking its C-terminal domain can still mature both the native diheme cyt *c* (NapB) and a heterologous monoheme cyt *c* (Cyt *c* C550). They contend that the C-terminal is not required for maturation because of functional redundancy between the two domains of CcmH. In contrast, Zheng et al. (Zheng et al., 2012) predicted the C-terminal of CcmH is a TRP (tetratricopeptide repeat) -like domain which recognizes apo-cyt *c* and concluded it is essential for cyt *c* expression. Our previous results suggest the C-terminal of CcmH is essential in expressing multi-heme heterologous cyt *c* (Su et al., 2020), agreeing more closely with Zheng et al. (Zheng et al., 2012).

To optimize the heme ligation process for Mtr cyt *c*, here we constructed a hybrid CcmH that uses protein domains from *E. coli* (the N-terminal of CcmH) and *S. oneidensis* MR-1 (the C-terminal of CcmI). We then explored the effects on cyt *c* expression and EET efficiency to test if this hybrid CcmH could boost the bioelectrical performance of Mtr-expressing *E. coli*.

## 2. METHODS

### 2.1 Construction of *E. coli* strains with *ccm* variants

We used Phusion DNA polymerase (NEB) for all the PCR amplification (please refer to Supplementary Information for cloning procedures). We have listed all the primers, plasmids, and strains we used in Table S1 and S2. We sequence-verified all plasmids used and transformed them into the *E. coli* C43(DE3) background (Lucigen) to construct all the strains.

### 2.2 Cyt *c* expression and growth conditions for *E. coli* strains

To express cyt *c*, we first inoculated all the *E. coli* strains directly from -80 °C glycerol stocks in 5 mL LB media with antibiotics (50 μg/mL kanamycin and 34 μg/mL chloramphenicol) and grew them at 37 °C overnight with 250 rpm sharking. Then we inoculated cell cultures (1:100 v/v ratio) in 50 mL 2xYT media in foil-covered 250 mL flasks with the same antibiotic concentrations and 1 mM δ-ALA (aminolevulinic acid), and grew these cultures at 37 °C with 250 rpm sharking. When the cell OD_600_ reached between 0.5 to 0.6, we added 4 μM IPTG to induce the expression of cyt *c*, followed by culturing at 30 °C with 250 rpm shaking overnight. Then we washed cells twice with M9 minimal medium (BD). All the chemicals and media are from Sigma-Aldrich unless otherwise noted.

### 2.3 Redness screening of the *E. coli* strains with *ccm* variants

We used redness screening for detecting the overall expression level of cyt *c*, as previously described (Su et al., 2020). We first grew all the strains from -80 °C glycerol stocks, then inoculated cell cultures in 2xYT media with foil-covered flasks (see **section 2.2**). When the cell OD_600_ reached between 0.5 to 0.6, we split the cell cultures into 96-well deep-well plates (Polypropylene 2.2 mL, VWR), where each well contained 0.5 mL cell culture. We then induced the cells with a gradient concentration of IPTG (0, 0.5, 1, 2, 4, 8, 16, 32, 62.5, 125, 250, and 500 μM). After overnight incubation, we transferred the cell cultures into 96-well white round-bottom microplates (3789A, Corning), and centrifuged the plate to pellet the cells. After removing the media, we used a scanner to take pictures of the plates and measured the redness of the cell pellets with a MATLAB image processing program (Su et al., 2020).

### 2.4 Enhanced chemiluminescence measurement of cyt *c* expression

We used enhanced chemiluminescence (ECL) to detect the cyt *c* expression as previously described (Su et al., 2020). In brief, we mixed 20 μL of 10 OD_600_ washed cells (described in **2.1**) with an equal volume of Laemmli Sample Buffer (2x), and heated the mixture to 95°C for 15 min. Then we loaded the samples and protein ladder (Precision Plus Protein WesternC Standards) into an SDS-gel (4-20% Crit TGX Stain-Free Gel) to perform SDS-PAGE (160 V, 45 min). Next, we transferred proteins from the SDS-gel onto the nitrocellulose membrane (Trans-Blot Turbo Midi NC Transfer Packs) via the Trans-Blot Turbo Transfer System (Bio-Rad). We then used ECL kit (Pierce Pico West Enhanced Chemiluminescence substrate, ThermoFisher) to perform the cyt *c* detection assay. Finally, we imaged the ECL signals using the FluorChem E system (ProteinSimple). All the chemicals and supplies are from Bio-Rad unless otherwise indicated.

### 2.5 Bioelectrochemical measurement

We used two-chamber, three-electrode electrochemical reactors to perform the electrochemical experiments as previously described (Su et al., 2020). The reactors contained three electrodes: a working electrode (graphite felt, Alfa Aesar) and a reference electrode (CHI111, Ag/AgCl with 3 M KCl, CH Instruments) in the anodic chamber, and a counter electrode (titanium wire, Alfa Aesar) in the cathodic chamber. We also used a cation exchange membrane (CMI-7000S, Membranes International) to separate these two chambers. Each chamber contained 150 mL M9 minimal medium (BD) and 50 mM D,L-lactate (Sigma-Aldrich) as the electrolyte. During the test, we continuously purged the anodic chamber with N_2_ gas to maintain anaerobic conditions, and we kept the temperature at 30 °C by placing all the reactors in an incubator. We used a potentiostat (VSP-300, Bio-Logic Science Instruments) to perform all the electrochemical measurements. We set the applied potential at 0.2 V versus the reference electrode to conduct chronoamperometry, and the electrical current (averaged at every 36 s) is reported as a function of the geometric surface area of the working electrode (0.0016 m^2^, two sides, diameter of 32 mm). For the differential pulse voltammetry (DPV) measurement (Xu et al., 2016), we scanned the potential from -0.4 V to 0.2 V versus the reference electrode, with the pulse height set to 50 mV, pulse width set to 500 ms, step height set to 1.0 mV, and step time set to 1000 ms.

## 3. RESULTS AND DISCUSSION

### 3.1 Design and construction of a hybrid cyt *c* maturation operon in *Escherichia coli*

To test our hypothesis that MR-1’s ligation module is more efficient at maturing MR-1 cyt *c* than *E. coli*’s ligation module, we made a series of modifications to the *ccmH* gene in the plasmid which carries the *ccm* operon from *E. coli* (**Figure 2**). We then co-transformed these plasmids with a separate plasmid that expresses *cymAmtrCAB* from MR-1 into *E. coli* C43(DE3) to create the wildtype *ccm, ccmHI, ccmH^N^,* and hybrid *ccmH* strains (**Table 1**). Specifically, to test the effect of altering the entire ligation module, we replaced *ccmH* from *E. coli* with *ccmHI* from MR-1, creating the I5101 plasmid in the *ccmHI* strain. To test the impact of replacing only the apo cyt *c* binding component, we fused the N-terminal region of the *E. coli ccmH* with the C terminal region of MR-1 *ccmI*, notated as the I5099 plasmid in hybrid *ccmH* strain. As a control, we also made the I5100 plasmid in *ccmH*^*N*^ strain from the I5099 plasmid, in which the C-terminal domain of *S. oneidensis* CcmI was removed. We also constructed a strain with the pACYC184 empty vector and *cymAmtrCAB* plasmid, dubbed the ‘no *ccm’* strain, as a negative control.

**Table 1.**
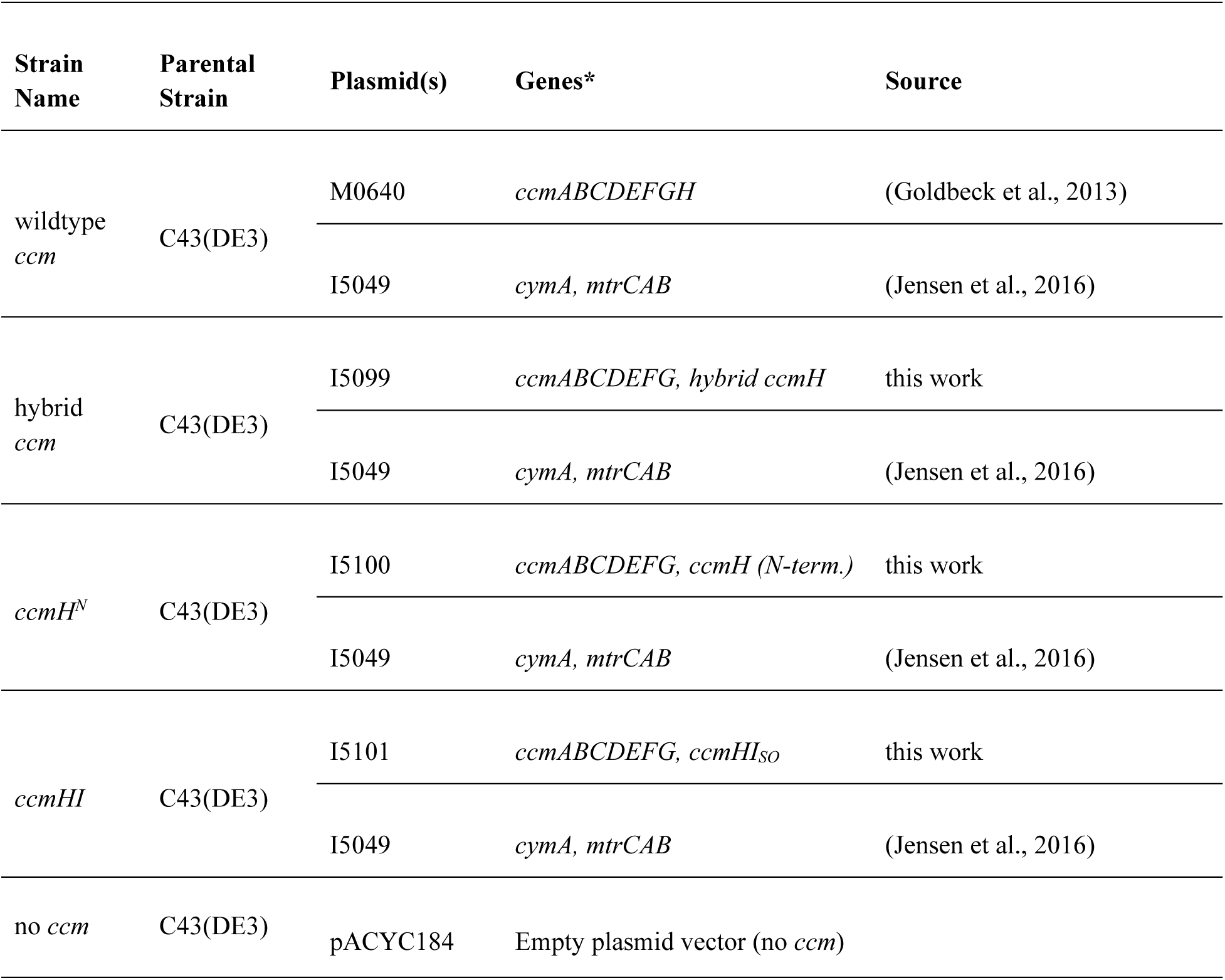
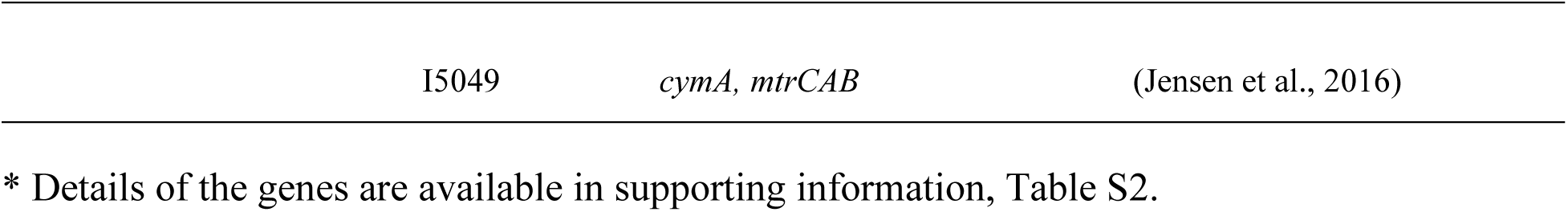
Strains Used in This Study.

**Figure 2.**
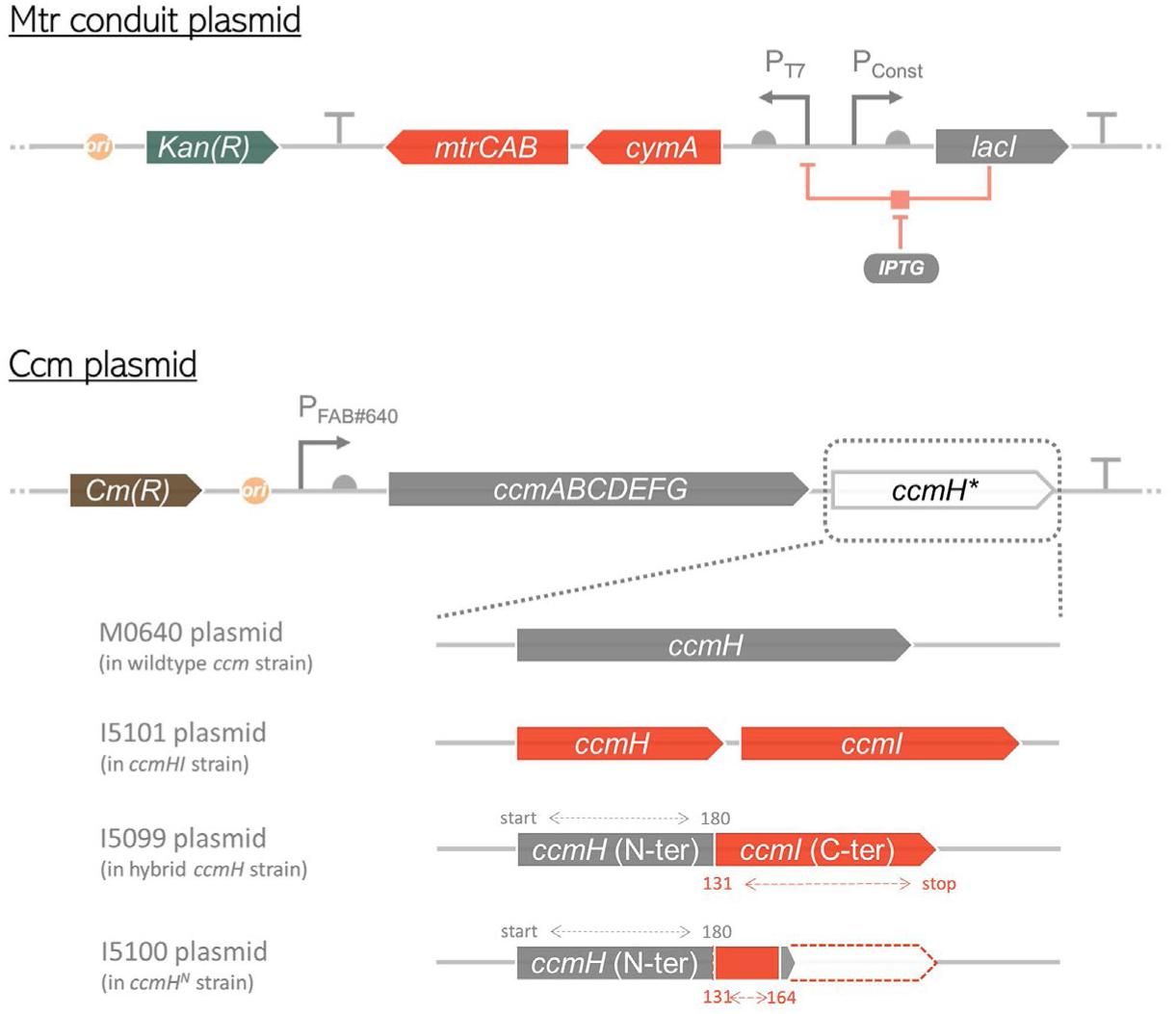
A schematic illustration describing the two plasmids we used for cytochrome *c* expression in engineered *E. coli*. For the Mtr conduit plasmid, *cymA, mtrC, mtrA, mtrB* are under the same T7 lacO promoter (P_T7_); for the Ccm plasmid, all the components are under a constitutive promoter (P_FAB#640_). In the Ccm plasmid, we replaced the coding sequence of *ccmH* from the *E. coli ccm* operon with three other variants. These plasmids coded for CcmHI from MR-1 (I5101), a fusion between the N terminal region of *E. coli ccmH* and the C terminal region of CcmI from MR-1 (I5099), and a control that removed the C terminal region of MR-1 CcmI (I5100). Gray and red regions indicate DNA regions that code for proteins from *E. coli* and MR-1, respectively. (origin of replication; promoter; ribosome-binding site, RBS; coding sequence, CDS; terminator).

### 3.2 The hybrid Ccm system can increase cyt *c* abundance in *E. coli*

To test if these *ccm* variants can mature and change the cyt *c* expression level, we first examined the cyt *c* expression of these *E. coli* strains with a redness assay (Su et al., 2020) across different IPTG induction levels (**Figure 3**). Since IPTG induction level is non-linear with protein expression, we present this data in term of relative promoter strength, which is the GFP expression per cell at a IPTG concentration normalized by the maximal GFP expression per cell (Goldbeck et al., 2013). The redness value, which indirectly measures the cyt *c* per cell, peaked at the relative promoter strength of ∼ 0.1 (arb. strength units) for all the strains with a Ccm plasmid. As expected, the no *ccm* strain has the lowest redness value because it cannot mature cyt *c,* and the wildtype *ccm* strain has a high redness value, consistent with its ability to mature cyt *c*. Compared to the wildtype *ccm* strain, the redness value of the *ccmHI* strain decreased by ∼40% (0.15 ± 0.003 arb. red units in *ccmHI* vs. 0.25 ± 0.003 arb. red units in wildtype). This decrease in cyt *c* expression might be due to a poor ability of the *E. coli* and MR-1 proteins to form a functional complex (Zheng et al., 2012). In contrast, the hybrid *ccmH* strain has a redness value (0.27 ± 0.002 arb. red units) that is 8% greater than the wildtype *ccm* strain, indicating that it can mature cyt *c* slightly better. This robust cyt *c* expression shows the C terminal region of CcmI from MR-1 can functionally replace the C terminal region of *E. coli* CcmH. Overall, these data show that the hybrid *ccm* system can increase cyt *c* abundance in *E. coli*, which potentially may increase its EET efficiency.

**Figure 3.**
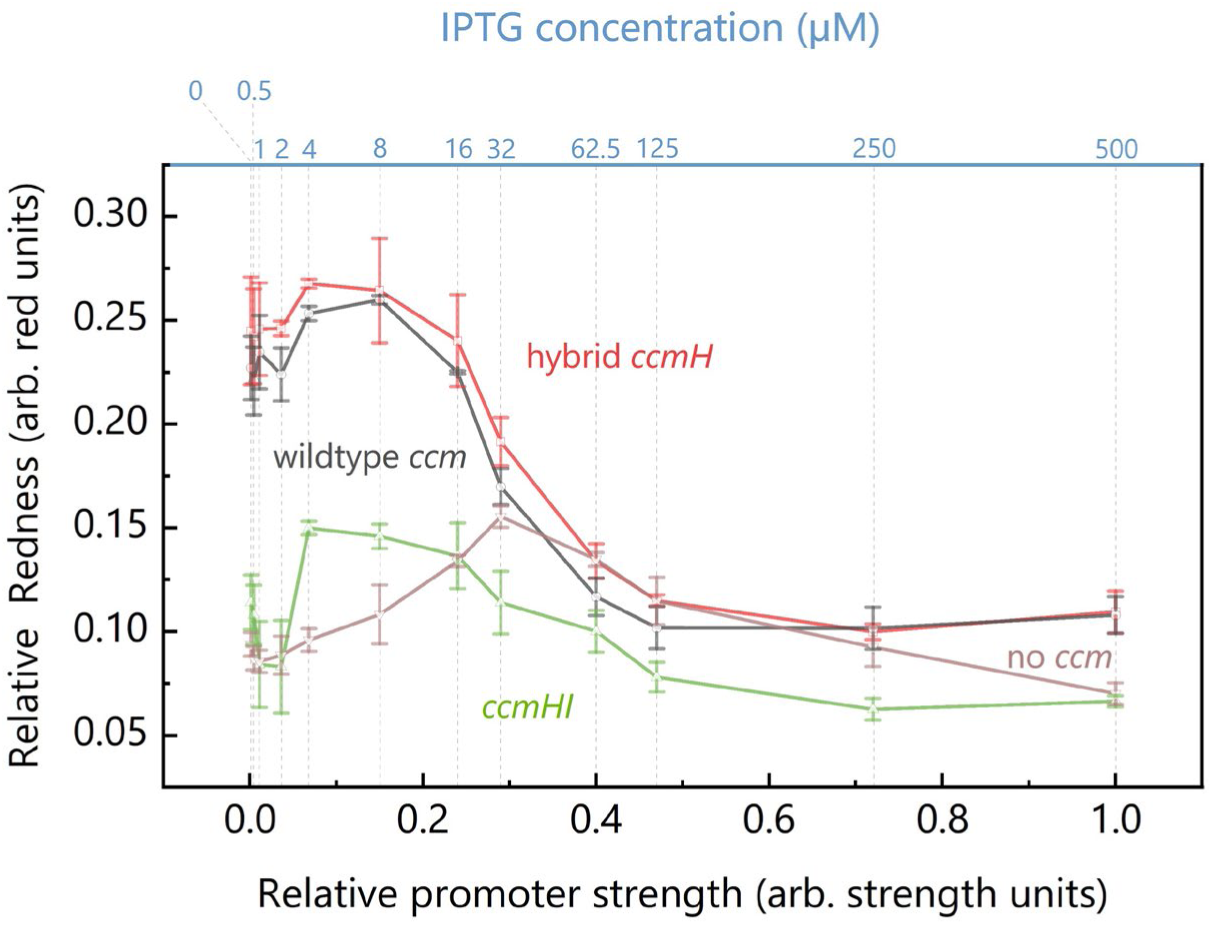
The hybrid *ccmH* strain expresses more cytochromes *c*. Relative redness as a function of relative promoter strength, i.e., induction level, for *E. coli* expressing *cymAmtrCAB* with different *ccm* operons. The hybrid *ccmH* strain has a higher level of overall cyt *c* expression than wildtype *ccm* strain. Results are representative of three independent experiments.

### 3.3 The hybrid Ccm system changes Mtr cyt *c* stoichiometry

To further investigate how the hybrid *ccm* system alters the expression of cyt *c* in the Mtr pathway, we used enhanced chemiluminescence (ECL) to characterize the abundance of CymA, MtrC, and MtrA in hybrid *ccmH* strain, wildtype *ccm* strain, and no *ccm* strain (**Figure S2**). These strains were all induced at 0.068 arb. strength units relative promoter strength (4 μM IPTG) since the hybrid *ccm* strain has one of the highest cyt *c* expressions at this induction (**Figure 3**). As expected, there was no detectable amount of cyt *c* in the no *ccm* strain (**Figure S2**). While the MtrC abundance was very similar, the abundance of MtrA decreased ∼15%, and CymA increased ∼60% in the hybrid *ccm* strain relative to the wildtype *ccm* strain (**Figure 4**). These data show that the hybrid *ccm* system increases the overall cyt *c* abundance by increasing the maturation of CymA.

**Figure 4.**
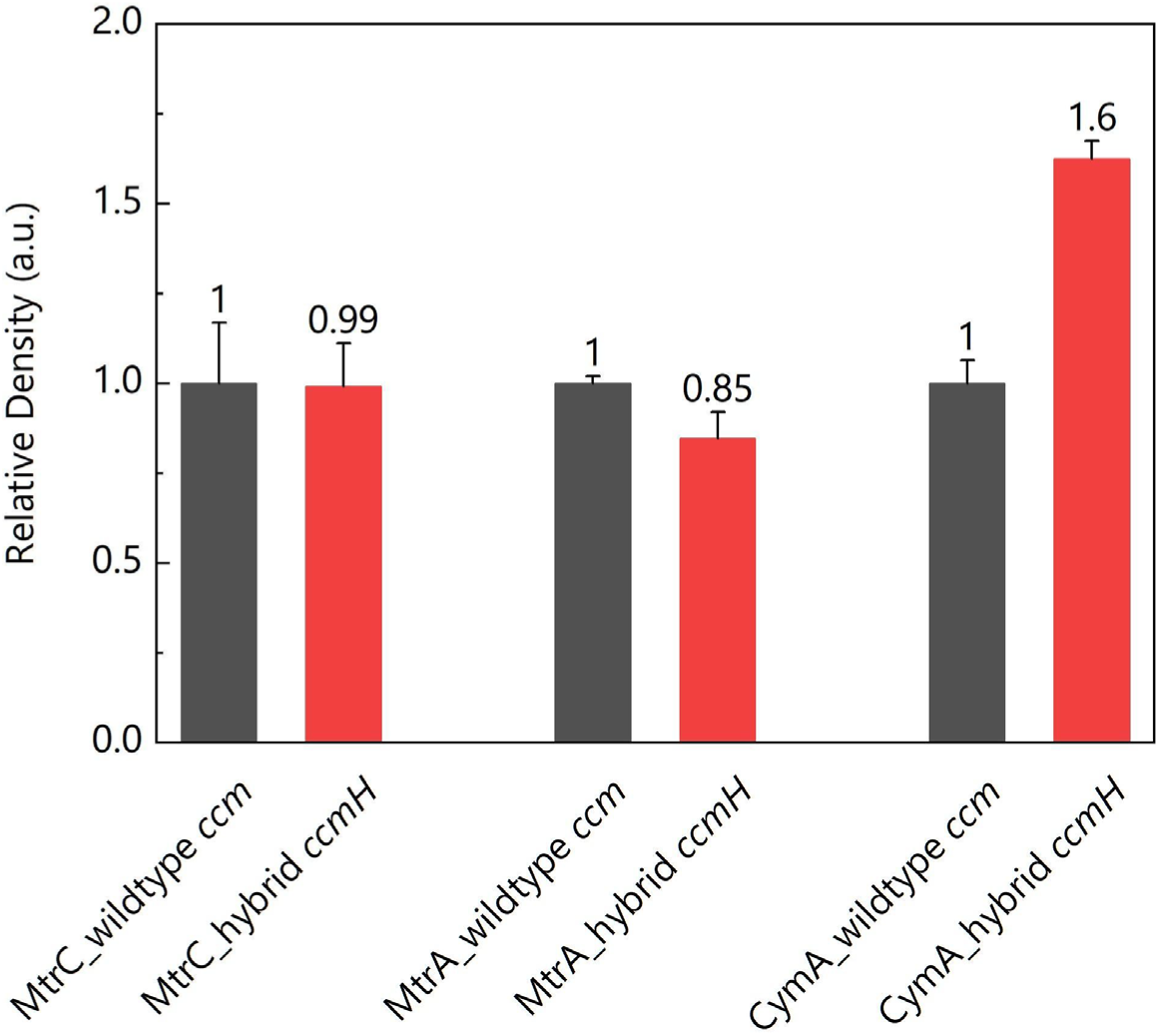
The hybrid *ccm* system increases the CymA expression in *E. coli*. The relative density of enhanced chemiluminescence (ECL) bands arising from three cyt *c* heterologously expressed in the hybrid *ccm* and wildtype *ccm* strains. The cyt *c* expression in negative control no *ccm* strain was below the limit of detection. Results are representative of three independent experiments.

### 3.4 The hybrid Ccm system increases the current generation of Mtr-expressing *E. coli*

Altering the stoichiometry of Mtr cyt *c* in *E. coli* can negatively or positively affect its bioelectrochemical performance (Su et al., 2020). To probe if the increase of CymA expression in hybrid *ccmH* strain will also enhance its bioelectrical performance, we measured the current generation (**Figure 5**) by the wildtype *ccm*, hybrid *ccmH, ccmH^N^* strains in three-electrode electrochemistry bioreactors. The wildtype *ccm* strain produced 11.47 ± 0.19 mA/m^2^, in line with our previous observations (Su et al., 2020). The hybrid *ccmH* strain produced 25.32 ± 3.61mA/m^2^, a 121% increase in current over the wildtype *ccm* strain. The *ccmH*^*N*^ strain produced only 3.42 ± 0.44 mA/m^2^, about a 70% decrease relative to the wildtype *ccm* strain, similar to the no *mtr* strain previously reported (TerAvest et al., 2014). Thus, we conclude that the hybrid *ccmH* strain matures more CymA and produces higher current than the *E. coli* Ccm system.

**Figure 5.**
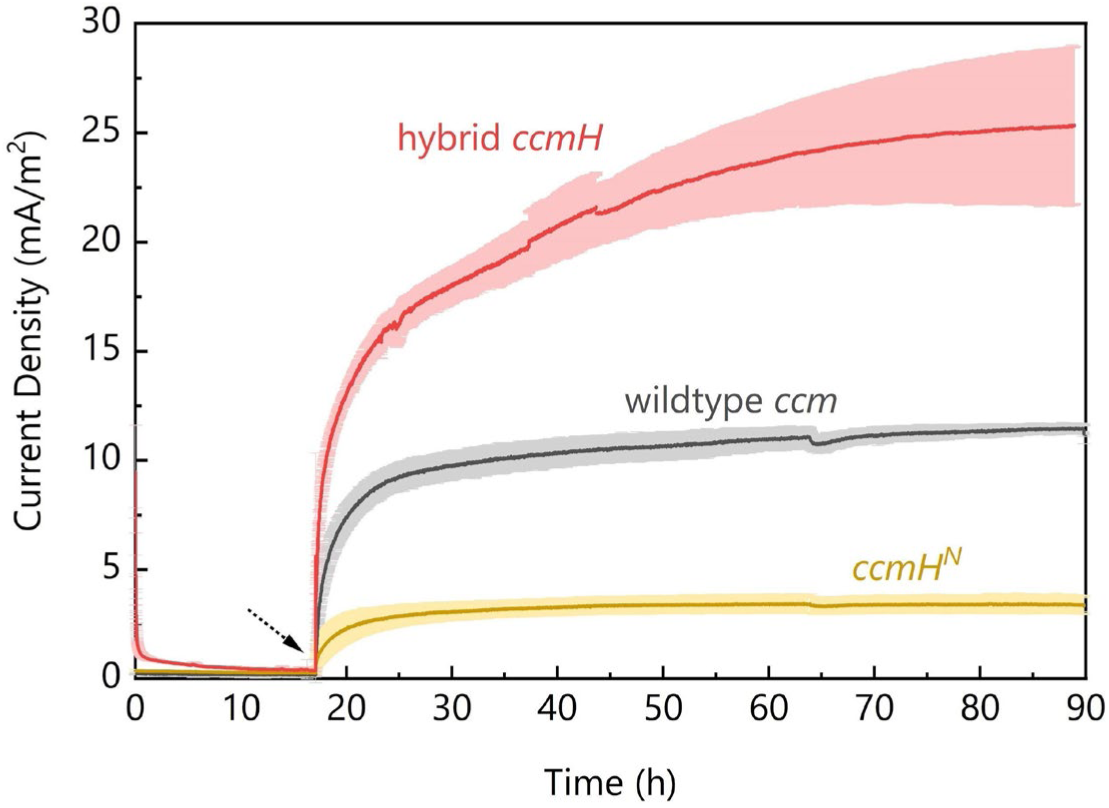
The hybrid *ccmH* strain produces a higher current. The current generation measured by chronoamperometry in the bioreactors showing hybrid *ccmH* strain produces more current than the wildtype *ccm* strain, while the negative control strain *ccmH*^*N*^ produces less current. Results are representative of three independent experiments. The dotted arrow shows the time the bacteria were injected.

To probe the mechanism underlying the increase in current production by the hybrid *ccmH* strain, we performed differential pulse voltammetry (DPV) (**Figure 6**) after strains reached the steady-state current generation after 90 h chronoamperometry testing. We observed a single anodic peak at redox potential (E_p_) of +51 mV_SHE_ in all three strains, which has been previously assigned to the MtrCAB complex (Okamoto et al., 2013). This observation confirms that all three strains use the MtrCAB conduit for the current production. The peak height in hybrid *ccmH* strain reached 8.31 μA, about 118% higher than the 3.82 μA in wildtype *ccm* strain, while in the *ccmH*^*N*^ strain, it was only 0.09 μA, about 2% of the wildtype. The ratio of the peak height for the hybrid *ccm* strain relative to the wildtype *ccm* strain (2.18) closely mirrors the ratio of steady-state current for these strains (2.21), indicating the increase in steady-state current arises from an increase in the flux of electrons through the MtrCAB complex. Thus, the hybrid *ccm* strain produces more current by more effectively maturing CymA, boosting its relative abundance, and increasing the electron flux through the MtrCAB conduit.

**Figure 6.**
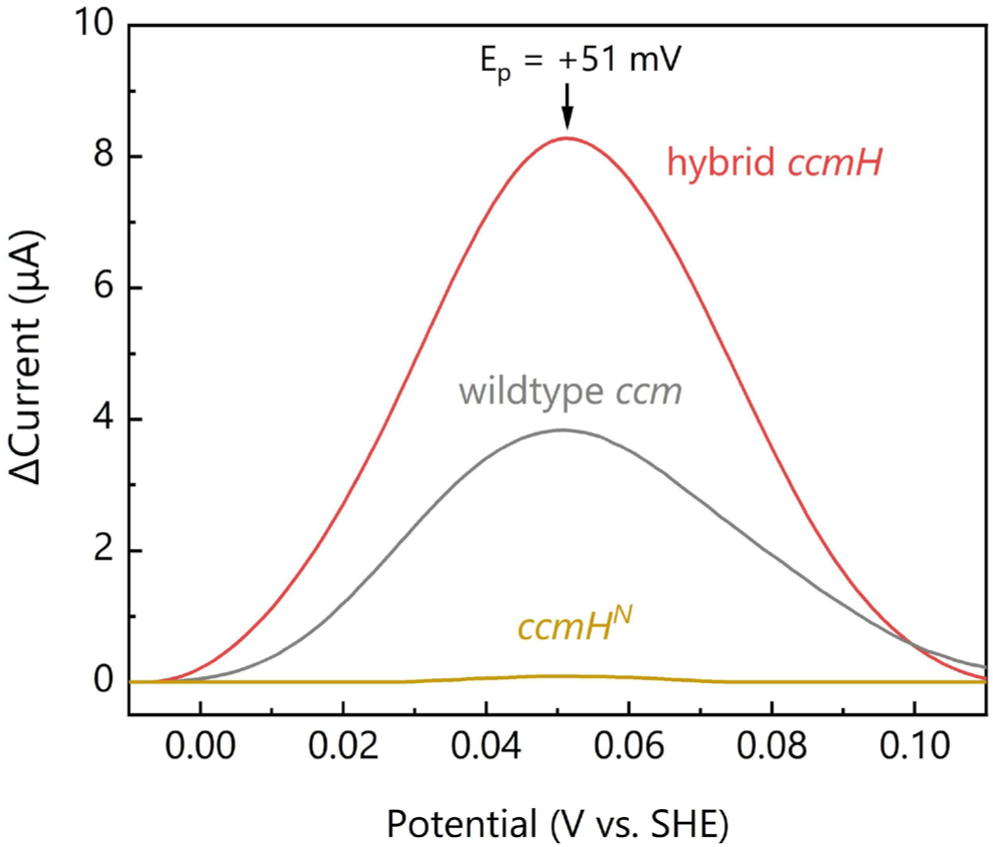
The hybrid *ccmH* strain shows higher redox activity through cyt *c*. Differential pulse voltammetry (DPV) shows that the hybrid *ccmH* strain has a higher redox peak at the cyt *c* related redox reaction potential (+51 mV vs. SHE). DPV signal smoothing and background deduction by QSoas Version 2.0 (Fourmond et al., 2009).

### 3.5 Hybrid *ccmH* strain produces more current per cell via both direct and indirect EET

While DPV confirms that the increase in electron flux through the Mtr pathway increases the current density, this increased flux could arise from an increase in the current per cell, an increase in the number of viable cells, or an increase in both. To distinguish between these possibilities, we measured both the current production and the number of viable cells using colony-forming unit counting in the *ccmH*^*N*^, hybrid *ccm*, and wildtype *ccm* strains (**Figure 7A**). The *ccmH*^*N*^ strain produced near baseline current on a per-cell basis (1.0 ± 0.005 fA·cell^-1^), as expected. Interestingly, the hybrid *ccmH* strains produced ∼50% more current per cell than the wildtype *ccm* (12.3 ± 0.3 fA·cell^-1^ versus 8.3 ± 0.4 fA·cell^-1^). These data confirm that the hybrid *ccmH* strain produces more current at the single-cell level by direct EET.

**Figure 7.**
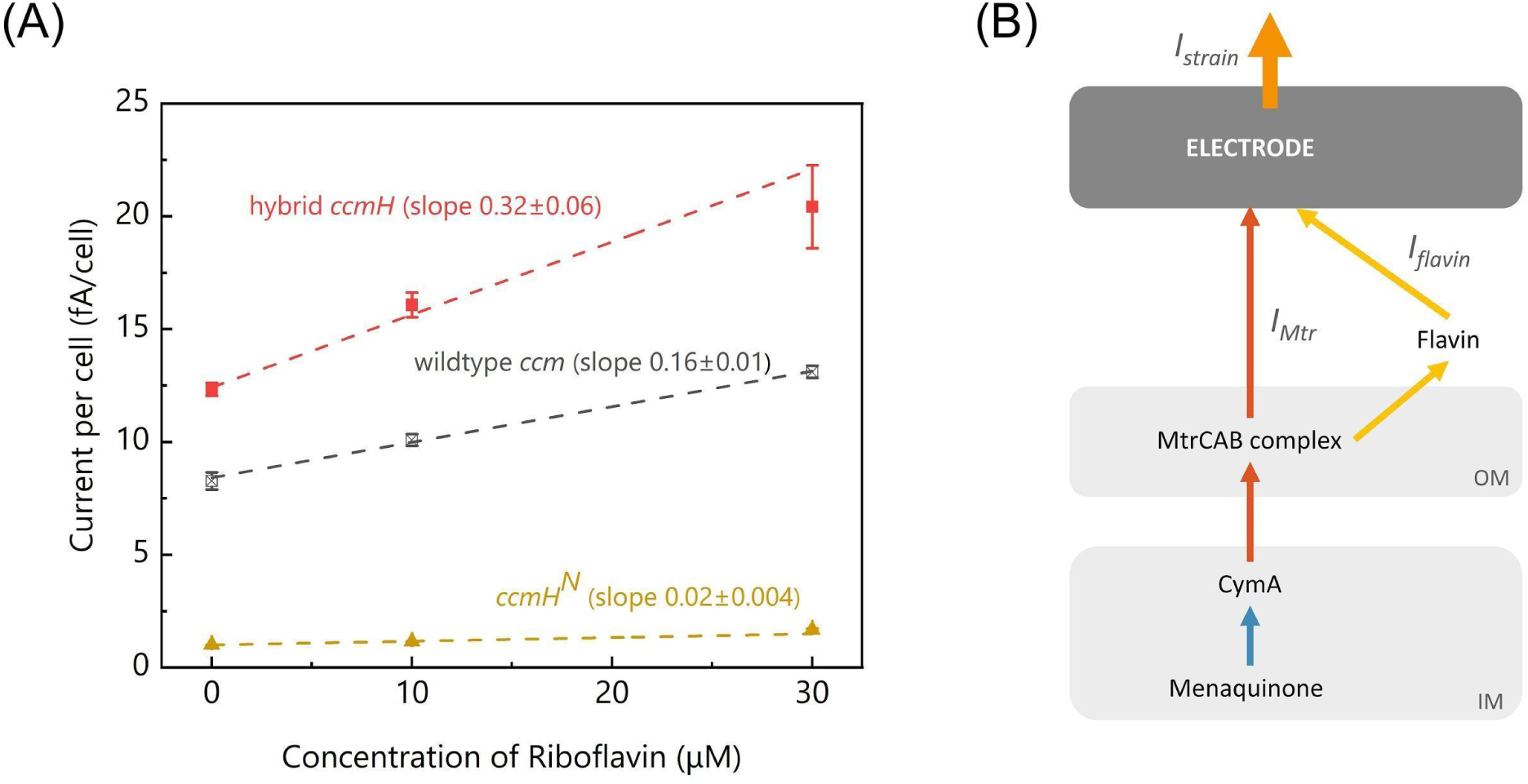
Current production by the hybrid *ccmH* strain is more sensitive to exogenous flavins. (A) The hybrid *ccmH* strain (red) has a more rapid and higher current response as a function of extra riboflavin concentration, compared to the wildtype *ccm* strain (gray), and *ccmH*^*N*^ strain (yellow). Results are representative of two independent experiments. Linear fit analysis (with Y error) is performed by OriginPro 2019b (OriginLab Corporation). (B) A schematic depicting the extracellular electron transfer via Mtr conduit. MtrCAB complex receives the electrons from CymA, then delivers to the electrode (*I*_*strain*_) through either MtrC (*I*_*Mtr*_) or flavin (*I*_*flavin*_). (OM: outer membranes, IM: Inner membranes).

In addition to direct EET via Mtr conduit, MR-1 (Brutinel and Gralnick, 2012; Coursolle et al., 2010) can also perform EET with flavins as a bound cofactor to cyt *c* (Edwards et al., 2015; Okamoto et al., 2014) or as a soluble mediator (Kotloski and Gralnick, 2013). To test if the hybrid *ccmH* strain can also utilize flavin for EET, we added exogenous flavins and measured the current density per cell. The addition of riboflavin to the *ccmH*^*N*^ strain resulted in a meager current increase (0.02 ± 0.004 fA·cell^-1^·µM^-1^). As expected, the current produced by both the wildtype *ccm* and hybrid *ccm* strains linearly increased with the exogenous flavin concentration (**Figure 7A**). However, the hybrid *ccm* strain produced significantly more current per µM exogenous flavin (0.32 ± 0.06 fA·cell^-1^· µM^-1^) than the wildtype *ccm* strain (0.16 ± 0.01 fA·cell^-1^·µM^-1^) (**Figure 7A**). These data indicate that the hybrid *ccm* strain produces more current at the single-cell level via a flavin-mediated EET as well as a direct EET mechanism.

### 3.6 Direct and flavin-mediated EET in the hybrid *ccmH* strain is limited by CymA

Our data suggest that the CymA amount limits electron flux and that the hybrid *ccmH* strain achieves higher current by enabling a greater electron flux from CymA into the same direct and indirect EET routes found in the wildtype *ccm* strain. To quantitatively test these hypotheses, we developed a quantitative model that relates EET in the hybrid *ccmH* strain to EET in the wildtype *ccm* strain and then tested whether this model could explain the current per cell measurement in the hybrid *ccmH* strain (**Figure 7**).

Explicitly, we modeled the overall EET current per cell (*I*_*strain*_) as the sum of the direct EET via Mtr conduit (*I*_*Mtr*_) and flavin-mediated EET (*I*_*flavin*_) (**Figure 7B**). We describe the flavin-mediated EET as the product of the flavin concentration ([*flavin*]) and a constant of proportionality (*k*_*strain*_), (**Equation 1**):

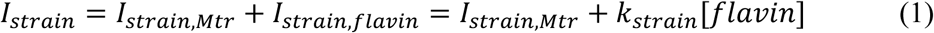

Assuming the amount of CymA is rate-limiting in our system (**Figure 4**), we can then calculate EET current in the hybrid *ccmH* strain (*I*_*hybrid*_) as a ratio of the CymA concentration in the two strains and the current in the wildtype *ccm* strain (*I*_*WT,Mt*r_) with the equation below (**Equation 2**):

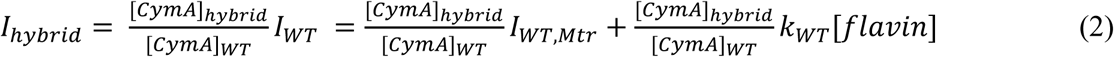

Thus, both the intercept and slope of the best-fit line for the hybrid *ccm* strain should be equal to the intercept and slope best-fit line of the wildtype *ccm* strain times the ratio of CymA concentration in these two strains.

Using the experimentally-determined values of *I*_*WT,Mtr*_ (8.3 ± 0.4 fA·cell^-1^), *k*_WT_ (0.16 ± 0.01 fA·cell^-1^·µM^-1^), and *[CymA]_hybrid_/[CymA]_WT_* = 1.6, we can predict the values of *I*_*hybrid*_ and compare it to our experimentally-determined values (**Figure 7A**). Both the predicted *I*_*hybrid, Mtr*_ and *I*_*hybrid, flavin*_ values, 13.28 ± 0.64 fA·cell^-1^ and 0.26 ± 0.02 fA·cell^-1^·µM^-1^, respectively, are quite close to our observation of the direct EET (12.3 ± 0.3 fA·cell^-1^) and mediated EET (0.32 ± 0.06 fA·cell^-1^·µM^-1^) in hybrid *ccmH* strain. This result provides strong evidence that the hybrid *ccmH* strain’s improved bioelectrochemical performance arises mainly from the CymA expression increase and underscores the importance of CymA in both direct EET and indirect EET.

## 4. DISCUSSION

In this study, we have shown that the C-terminal region of CcmH can be complemented by the homologous region of CcmI from *S. oneidensis* MR-1, which increases the bioelectronic performance of Mtr-expressing *E. coli*. We found that the hybrid CcmH can increase the overall cyt *c* expression (**Figure 3**), with a preferred 60% increase in CymA expression (**Figure 4**). This increase in CymA expression led to a 48% more EET current per bacterium and doubled the electrical response to exogenous flavin (**Figure 7**), consistent with CymA being rate-limiting in EET. In the following, we discuss how these findings impact the development of biofuel cells and biosensors.

First, our data show that the C-terminal region of CcmH is essential for cyt *c-*based EET. *Shewanella* and *Geobacter* species are the dominant organisms used for EET applications, such as MFCs. These microorganisms express a significant amount of cyt *c* to perform EET (Meyer et al., 2004; Shi et al., 2009). As a result, there are increasing efforts focused on genetically introducing cyt *c* to make new organisms capable of EET (Schuergers et al., 2017; Sekar et al., 2016; Thirumurthy and Jones, 2020), overexpressing cyt *c* to understand the structural basis of EET (Edwards et al., 2020, 2019; Wang et al., 2019), and making cyt *c* -based conductive biomaterials for bioelectrical applications (Liu et al., 2020; Sun et al., 2018). Previously we used a random mutagenesis approach to improve cyt *c* maturation, which yielded strains that both increased current by up to 77% and decreased current production by as much as -66% (Su et al., 2020). Leveraging understanding from our previous study, this work presents a rational approach that increased current by 121%. Thus, understanding how cyt *c* mature can provide guidance on manipulating cyt *c* expression, which will assist both fundamental studies and these applied efforts to increase the bioelectrical performance of MFCs.

Second, we found that the hybrid *ccmH* strain generates higher electrical signals using both Mtr and flavin-mediated EET. Increasing the EET efficiency of Mtr-expressing *E. coli* is a particular interest for biosensing because it can increase the signal-to-noise ratio, enabling higher sensitivity detection. In recent years, much progress has been made in engineering the electrical communication between bacteria and electrodes, such as responding to specific chemicals using chemical-sensitive promoters (Golitsch et al., 2013; Webster et al., 2014; Yang et al., 2018), miniaturized reactors with optimized structure design (Zhou et al., 2017), and modified material-bacteria interface (Zajdel et al., 2018). Enhancing the bioelectrical performance of Mtr-expression *E. coli* offers a reliable chassis in biosensing applications.

Lastly, our work provides guidance us on how to build a more efficient EET conduit for bioelectrical sensing. Our model for the current generation in hybrid *ccmH* strain together with the fact that CymA can only carry 40% (fewer heme groups per protein) as many electrons as MtrA or MtrC, strongly suggests that electron transfer through CymA is a bottleneck for EET in the Mtr-expression *E. coli*. Future work will focus on increasing the electron flux between menaquinone and the MtrCAB complex to overcome this rate-limiting step. Also, since the electrical response is linear in both CymA amount and flavin concentration, this informs more information rich sensing scenarios, including building an “AND” gate with dual inputs.

## 5. CONCLUSIONS

In summary, a hybrid Ccm system altered the stoichiometry of the expressed cyt *c,* thus accelerating the rate-limiting step in EET and enhancing the bioelectronic performance of *E. coli*. Our results suggest that increasing the electron flux from menaquinone to MtrCAB complex could further enhance the bioelectrical performance of engineered *E. coli*. More broadly, our work provides a new strategy to express cyt *c* that underlies microbial bioelectronics and opens new possibilities for engineering more sophisticated and sensitive biosensors.

## SUPPLEMENTARY DATA

Supporting Information is available online.

## AUTHOR CONTRIBUTIONS

L.S. contributed to strain construction, design of the study, conducting experiments, analysis of the data, and writing and revision of the manuscript. T.F. contributed to strain construction, design of the study, conducting experiments, analysis of the data, review and revision of the manuscript. C.M.A-F. contributed to the design of the study, analysis of the data, and writing of the manuscript.

## ACKNOWLEDGMENT

We thank Dr. Shuai Xu and Prof. Mohamed Y. El-Naggar, the University of Southern California for discussions on the DPV experiments. Work at the Molecular Foundry was supported by the Office of Science, Office of Basic Energy Sciences, of the U.S. Department of Energy [DE-AC02-05CH11231]. This work was supported by the Office of Naval Research, [N0001418IP00037]; and the China Scholarship Council [201606090098 to L.S.].

