## Supplementary Information for "A hybrid cyt *c* maturation system enhances the bioelectrical performance of engineered *Escherichia coli* by improving the rate-limiting step"

### Supporting Methods

**Construction of I5101 plasmid for expressing *E. coli* CcmABCDEFG and *S. oneidensis* CcmH and CcmI.** For *ccmHI* plasmid construction *S. oneidensis ccmH*, *ccmI* and upstream region of *ccmI* (CAAACTAAAGTGAATATAGAAACA) were amplified by PCR using *S. oneidensis* MR-1 chromosome as a template and Shew-CcmHgib-F3 and Shew-CcmHgib-R primers for *ccmH* amplification and Shew-CcmIgib-F and Shew-CcmIgib-R3 primers for *ccmI* amplification. An entire plasmid of M0640 (*ccm*) except *E. coli ccmH* was amplified by PCR using M0640-CcmG-R2 and M0640-CcmHout-F3 primers. The amplified three fragments were assembled using a Gibson Assembly Master Mix (New England BioLabs) and *E. coli* Mach1 was transformed using the sample to construct *ccmHI* plasmid. The ribosome binding site (RBS) of *ccmH* is within the *E. coli ccmG* gene (AAGGAGG) and the possible RBS of *ccmI* is located in the upstream region although the site seems to be unclear.

**Construction of I5099 plasmid for expressing hybrid CcmH (fusion of N-terminal domain of *E. coli* CcmH and C-terminal domain of *S. oneidensis* CcmI).** The C-terminal domain of *S. oneidensis ccmI* (131 aa to the end) that is a TPR (tetratricopeptide repeat) domain was amplified by PCR using CcmI-Cter-F and Shew-CcmIgib-R3 primers and *S. oneidensis* MR-1 chromosome as a template. The amplified fragment was purified by gel extraction and then used as a primer to amplify the entire plasmid of M0640 (*ccm*) by PCR. The amplified fragment containing *E. coli ccmABCDEFG* and hybrid *ccmH* (N-terminal domain of *E. coli* CcmH [1 aa to 180 aa] and C-terminal domain of *S. oneidensis* CcmI) was ligated and *E. coli* Mach1 was transformed using the ligated sample to construct hybrid *ccmH* plasmid.

**Construction of I5100 plasmid for expressing the N-terminal domain of *E. coli* CcmH.** To confirm that the C-terminal domain of *S. oneidensis* CcmI is very important for the multiheme cytochrome *c* production and for improvement of electron transfer rate, the *ccmH*$^{N}$ plasmid was constructed. The hybrid *ccmH* plasmid was digested with HindIII and the digestion site of the plasmid was blunted by T4 DNA polymerase to create a stop codon. The plasmid was self-ligated and then transform *E. coli* DHα with the ligated solution, resulting in *ccmH*$^{N}$ plasmid. The plasmid contains the fusion gene of the N-terminal of the *E. coli* CcmH domain (1-180 aa) and *S. oneidensis* (131-164 aa [AANPHAGMDTAQIMAQRVQMMEAQVQAEPENSQA]) and SFDR peptide (*E. coli* CcmH domain fused with $A_{131}$ANPHAGMDTAQIMAQRVQMMEAQVQAEPENSQ$A_{164}$SFDR). The CcmH$^{N}$ does not contain the TPR domain.

**Table S1. Primers used in this study**

| **Primer name** | **Sequence (5´→3´)** |
|---|---|
| Shew-CcmHgib-F3 | ctgtgggagaaatacagtaaggaggccgcacaATGAGAACACTGACAAAAATCATCG |
| Shew-CcmHgib-R | ctatattcactttagtttgTCATTTACTGTCACGCTTCAATAAG |
| Shew-CcmIgib-F | CAAACTAAAGTGAATATAGAAACAATG |
| Shew-CcmIgib-R3 | cataaactaagcgcactttgtcaacagtctagccgatTTATTGTACTTGAGTATCCAGTACTA |
| M0640-CcmG-R2 | TGTGCGGCCTCCTTACTGTATTTCTCCCACAGC |
| M0640-CcmHout-F3 | GGCTAGACTGTTGACAAAGTGCGCTTAGTTTATG |
| CcmI-Cter-F | ggcaattatcagcaggtgaaaatctggcagcaggccGCTGCTAACCCGCATGCAGGTATG |

The small letters are an additional sequence.
*The letters with a same color are the homologous region that is used for the Gibson assembly plasmid construction.

**Table S2. Genes used in this study**

| Gene | Organism | Gene ID | Uniprot ID |
|---|---|---|---|
| *ccmA* | *Escherichia coli* K12 | 946714 | P33931 |
| *ccmB* | *Escherichia coli* K12 | 946692 | P0ABL8 |
| *ccmC* | *Escherichia coli* K12 | 946703 | P0ABM1 |
| *ccmD* | *Escherichia coli* K12 | 946709 | P0ABM5 |
| *ccmE* | *Escherichia coli* K12 | 946697 | P69490 |
| *ccmF* | *Escherichia coli* K12 | 948783 | P33927 |
| *ccmG* | *Escherichia coli* K12 | 949073 | P0AA86 |
| *ccmH* | *Escherichia coli* K12 | 946623 | P0ABM9 |
| *cymA* | *Shewanella oneidensis* MR-1 | 1172176 | Q8E8S0 |
| *mtrC* | *Shewanella oneidensis* MR-1 | 1169552 | Q8EG34 |
| *mtrA* | *Shewanella oneidensis* MR-1 | 1169551 | Q8EG35 |
| *mtrB* | *Shewanella oneidensis* MR-1 | 1169550 | Q8CVD4 |
| *ccmH* | *Shewanella oneidensis* MR-1 | 1168152 | Q8EK35 |
| *ccmI* | *Shewanella oneidensis* MR-1 | 1168149 | Q8EK38 |

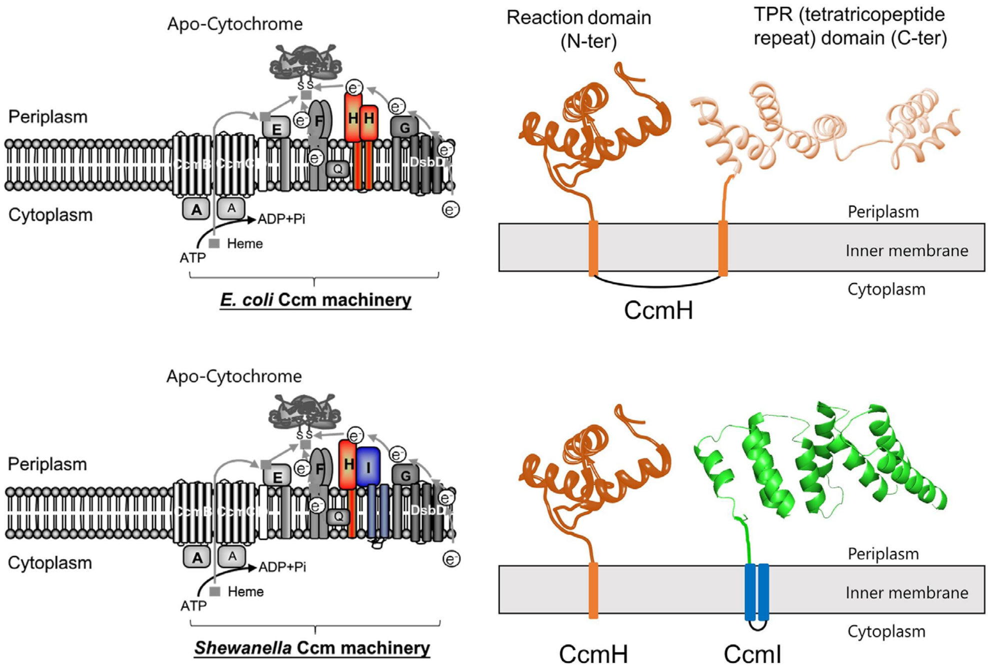


**Figure S1.** A schematic illustration showing the difference between the Ccm systems in *E. coli* and *Shewanella*. CcmH in E. coli is divided into CcmH (with the reaction domain can perform ligation) and CcmI (with the TPR domain to bind apo-cyt *c*) in *S. oneidensis*.

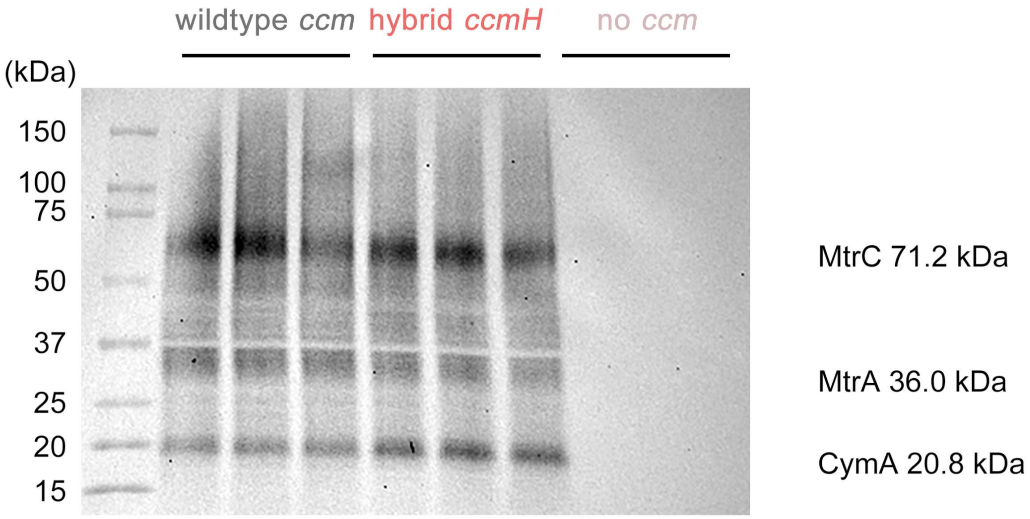


**Figure S2.** Enhanced chemiluminescence (ECL) analysis reveals the cyt *c* expression (MtrC, MtrA, and CymA) changes between wildtype *ccm* strain, hybrid *ccmH* strain, and no *ccm* strain.
